# Virus infection significantly decreases insect fitness: a meta-analysis

**DOI:** 10.1101/2025.01.19.633787

**Authors:** Cássia Siqueira Cesar, Vitória Horvath Miranda, Elielson Rodrigo Silveira, Thiago Assis de Oliveira, Rodrigo Cogni

**Affiliations:** Department of Ecology, University of São Paulo, São Paulo, Brazil; Department of Botany, University of São Paulo, São Paulo, Brazil

**Keywords:** ecological interactions, host-pathogen, parasitism, pathogens, vector

## Abstract

Organisms are constantly at risk of being infected by pathogens such as viruses. Adaptations against viral infection include immune defenses encoded within the host genome and associations with defensive symbionts such as *Wolbachia. Wolbachia* is widely spread among insects, and its success in nature may be due to its antiviral effects, which can benefit hosts if viruses significantly reduce host fitness. Here, we conducted a meta-analysis to assess the degree to which viral infection affects the fitness of insect hosts, and which factors may influence the impact of viral infection on hosts such as if the insect host is a vector or not of viruses that cause diseases in humans and plants, and if the insect is a new or a natural host of a specific virus. We gathered 1,040 effect sizes from 150 studies. Our results show that viruses significantly reduce host fitness, especially their survival. The decrease in host fitness is higher in non-vector than in vector insects, and we found no difference in fitness decrease caused by viral infection between new and natural hosts. Moreover, we found that fitness effects caused by viruses vary between host order and fitness components. In conclusion, our results show that viruses exert severe harmful effects on hosts by decreasing their fitness. In this context, harboring symbionts that confer antiviral protection, such as *Wolbachia*, can be highly advantageous for hosts, enhancing their fitness. Conversely, *Wolbachia* can benefit from the presence of viruses, facilitating its spread in insect populations by offering antiviral protection.

## Introduction

Organisms are constantly at risk of being infected by a great variety of microorganisms in nature, including pathogens and parasites such as viruses, bacteria, fungi, nematodes, and protozoans [1, 2]. Because pathogens can affect the fitness of hosts by decreasing their longevity and fecundity, pathogens can exert selective pressure for hosts to evolve resistance mechanisms against infections [3], and in response, pathogens can counter-adapt the resistance mechanisms that hosts have evolved [4]. Adaptations to resist pathogens include inherent immune defenses encoded within the host genome and associations with symbionts that protect hosts against infection [5]. For example, the *pastrel* gene in *Drosophila melanogaster* (Diptera) is associated with resistance against *Drosophila* C Virus (DCV) [5, 6], but flies can also be associated with bacterial symbiont *Wolbachia* that confers protection against a wide range of RNA viruses [7–10]. *Wolbachia* is commonly found infecting wild insect populations, and its mutualistic effect of protecting hosts against viral infection may be one of the factors for this symbiont being so well distributed among insect hosts. However, given that the association with certain *Wolbachia* strains can be costly for hosts, this association would be advantageous to hosts only if the viruses infecting them cause a significant reduction in their fitness. Therefore, *Wolbachia’s* antiviral effect could help explain its success in nature only if the association between hosts and viruses is harmful for hosts.

Studies on insect-virus association have been increasing in number in the past decades, given its importance to the study of host-pathogen evolution. Many studies focus on understanding the interaction between insects and viruses that are pathogenic and potentially lethal, such as RNA viruses in *Drosophila* species [11–14] and baculoviruses in Lepidoptera [15–17], the letter being of great importance in the field of biological control as some of these viruses are used as biocontrol agents for crop pests [17]. Traditionally, to study viruses, researchers have methods to isolate and replicate them in the laboratory, for example using cell culture. Such methods have helped to gather most of the current knowledge we have on viruses. However, some viruses are difficult to grow in the laboratory, restricting our knowledge [18, 19]. Recent advances in molecular biology allowed studies of metagenomics to discover many new viral species [20]. For example, a recent study identified 30 different viruses in a single population of *D. melanogaster*, including 13 viruses that had not previously been associated with this species [10]. In another study on *Apis mellifera* (Hymenoptera), researchers found 30 distinct viruses infecting the colonies, most of which had never been detected in honeybees before [21]. Although there is a widely accepted notion that viruses are always pathogenic [22], the effects of recent-discovered viruses on host health remain largely unknown [23]. Therefore, although recent studies on the insect-virus association show that hosts can be infected by a great variety of viruses, a question remains to whether most of these interactions are harmful (i.e., parasitism) or neutral (i.e., commensalism). Moreover, the advance in the field not only helps with the discovery of new viruses of ecological importance for wild insect populations, but it also contributes for the discovery of new viruses that can potentially cause human and plant diseases, such as viruses carried by insect vectors.

Understanding the interaction between viruses and their insect vectors is of great importance to the field of public health as it includes research on the evolution and transmission of arboviruses such as Dengue and Chikungunya [24, 25], and to the agriculture field as it includes research on plant viruses that cause great loss in crops such as tomato yellow leaf curl virus and rice stripe virus [26, 27]. Viruses that infect insect vectors are a particular case, as they need to adapt to replicate in two distinct hosts [24, 28], and they depend on the survival of their vector to complete their cycle and be transmitted to their next host [15, 29]. Therefore, because efficient virus transmission depends on balancing the virus infection and the health of the insect vector [27], we expect the negative impact of viruses on the fitness of their vectors to be less significant compared to non-vector insects. Moreover, these studies contribute to understanding how infectious diseases can arise and how host-virus dynamic works in nature.

The host-virus dynamic depends on host and viral effects. Host effects include susceptibility to new infections, which depends on defense mechanisms such as resistance (i.e., when hosts can control and reduce viral load due to the activation of their immune system) and tolerance (i.e., when hosts reduce the impact of the infection on their fitness without reducing viral load) [30, 31]. Viral effects include virulence (i.e., the level of harm that the virus causes to a host) [31, 32]. These traits are related to host shifts and the adaptation of viruses to new and natural hosts [31, 32]. Susceptibility and virulence are expected to be higher in new hosts, especially if the virus jumps between distantly related hosts [31–34]. Therefore, the harm that a virus can exert on host fitness may depend on whether the host species is new or natural. New hosts are expected to have higher mortality rates and greater reduction in their fitness in general than natural hosts.

Although there are numerous studies on viral infection in insects, the overall extent of the negative impact that viruses have on the fitness of their hosts remains unclear. Here, we conducted a meta-analysis that aimed in reviewing the current knowledge we have on viral infection in insects, quantifying the degree to which viral infection affects host fitness, and which factors may influence the impact of viral infection on hosts. Specifically, we asked: 1) what is the magnitude of effect of the harm that viruses have on the fitness of their hosts, 2) if the negative effect of the virus on the fitness of hosts is stronger in vector than in non-vector insects, and 3) if the negative effect of the virus on fitness is stronger in new hosts than in natural hosts. Moreover, we tested if the magnitude of effect differed between host orders, fitness components, and different methodological approaches.

## Material and Methods

### Literature search

We conducted a literature search using all databases of ISI *Web of Science* and *Scopus* platforms. We used the following combinations of keywords: “(insect* OR hymenoptera OR coleoptera OR lepidoptera OR trichoptera OR blattodea OR orthoptera OR diptera OR odonata OR dermaptera OR siphonaptera OR diptera OR mantophasmatodea OR hemiptera OR homoptera OR grylloblattidae OR neuroptera OR phthiraptera OR mantodea OR ephemeroptera OR megaloptera OR psocoptera OR mecoptera OR plecoptera OR strepsiptera OR isoptera OR thysanoptera OR heteroptera OR phasmida OR embioptera OR apterygota OR fly OR flies OR drosophila OR mosquito OR mosquitoes OR aedes OR culex) AND (infec*) AND (virus OR viruses OR viral) AND (fitness OR fecundity OR fertility OR “egg hatch” OR survival OR longevity OR lifespan OR “developmen* time” OR “body size”)”. We found a total of 3,752 abstracts to read.

### Inclusion criteria

For the title and abstract screening, study must have: (1) been performed on insect hosts, (2) mentioned that the host was infected with virus, (3) mentioned any fitness measure such as survival, longevity, lifespan, fecundity, fertility, egg hatch, development time, body size, etc. After abstract screening, our sample had 506 potential studies to be included.

For the full text screening, study must have: (1) been written in English, Portuguese or Spanish, (2) been empirical (reviews and studies with mathematical modelling were excluded), (3) measured any fitness measure (i.e., response variable) of the hosts, such as survival, longevity, lifespan, fecundity, fertility, egg hatch, development time, body size, etc., (4) compared treatment (host infected with virus) and control (host uninfected with virus), and (5) reported sample size, mean and standard deviation (S.D.) or standard error (S.E.), proportion or number of events of the response variables. If descriptive data was missing, the study must have reported inferential statistics (F, t, and degree of freedom) to be included. (6) In the case of studies performed in insects that are vectors of viruses that infects plants, we excluded all studies in which the insects were reared most of their lifetime in infected/sick plants as any effect on the fitness of these insects could be either because of the direct viral infection or because the virus changed host plant traits such as chemical defense levels or nutritional quality. Studies in which insects were exposed to infected plants for a short period of time just to acquire the virus and then reared in healthy plants, were included in our analysis. (7) If insects were reared in infected hosts, such as parasitoids reared in larvae infected with virus, we excluded the study from our sample as any effect on the fitness of the insect feeding on infected hosts for most of their lifetime could be because of the direct effect of viral infection or because the infected host have different nutritional qualities. After full text screening, our sample included 170 studies.

### Data extraction

We extracted different fitness measures and grouped similar data types under four broad categories of insect performance: body size, development time, fecundity, and survival (Table S2). When possible, we extracted the raw data from tables and graphs. Otherwise, we collected data from inferential statistics (F, t, and degree of freedom). When data was reported as mean ± S.E., we calculated the S.D. based on the S.E.. In the case of missing S.D. or S.E., we estimated the S.D. using the metric proposed by Bracken (1992) [35]. When data was reported in figures, we extracted it using *PlotDigitizer* [36]. In the case of data reported in box plots, we estimated the mean and S.D. using the metric proposed by Hozo et al. (2005) [37]. If the figure reported was a survival curve, we extracted the median and calculated the estimated S.D., using the Bracken (1992) metric [35]. We considered the median equal to the mean when the sample size was ≥ 25 [37].

Besides the fitness measures as response variables to calculate effect sizes, we collected aspects of the original studies that could be used as moderators in our models, such as: 1) vector: if the insect is a vector for viruses that causes diseases in animals or plants or if it is a non-vector, 2) host order: insect order that the host belongs to, 3) fitness component: the four broad categories of fitness measure (body size, development time, fecundity, and survival), 4) host type: if the insect is a natural or a new host of a specific virus, 5) virus type: if it is a DNA or RNA virus, 6) virus inoculation: method used to inoculate the virus in hosts; 7) life stage: life stage of the host in which the viral effect was measured (Table S3).

### Effect size calculation

When data was reported as proportion or number of events, we used the odds ratio (OR) to calculate effect sizes. The odds ratio estimates the chance of an event occurring in one group relative to chance of the same event occurring in the other group [38]. All the data we collected were related to survival, either as proportion or number of events. Therefore, we compared the odds of survival in hosts infected with the virus to the odds of survival in hosts not infected. For data reported as mean ± S.D., we used the Hedge’s g to calculate the effect sizes. Hedge’s g is the standardized mean difference between treatment and control groups corrected for bias [39]. When dealing with different estimates of effect size in a dataset, it is necessary to convert all effect sizes into a common metric, therefore we converted the OR estimates to Hedge’s g [40]. The calculation of effect sizes, including the conversion of OR to Hedge’s g, was performed using the function *escalc* in the *metafor* package in R [41, 42]. To calculate Hedges’ g based on inferential statistics, we used the Practical Meta-Analysis Effect Size Calculator [43], which uses pre-established equations to estimate effect sizes from different inferential statistics.

We calculated and adjusted the direction of effect sizes in a way that negative values of Hedges’ g indicate that viruses have a negative effect on the fitness of hosts. Therefore, negative values mean decreased fecundity, survival, and body size, but increased development time. Positive values indicate that viruses had a positive effect on host fitness.

### Statistical analysis

To account for data non-independence [44], we built multilevel meta-analytical models with four random effects: study ID, effect size ID, host phylogeny and host species. We obtained the insect phylogeny from Chesters et al. (2020) [45]. We removed from our dataset studies with species that were not found in the insect phylogeny (N=16 species) and then pruned the insect tree according to the species included in our dataset. We did not include virus phylogeny in the analysis due to the unavailability of comprehensive phylogenetic data on viruses. Our final dataset included 150 studies, 55 host species (Fig 1), and 90 study systems (Table S1). The full screening process following the PRISMA protocol [46] (Fig S1), and the list with all studies included in the meta-analysis are in the supplementary information.

**Fig 1.**
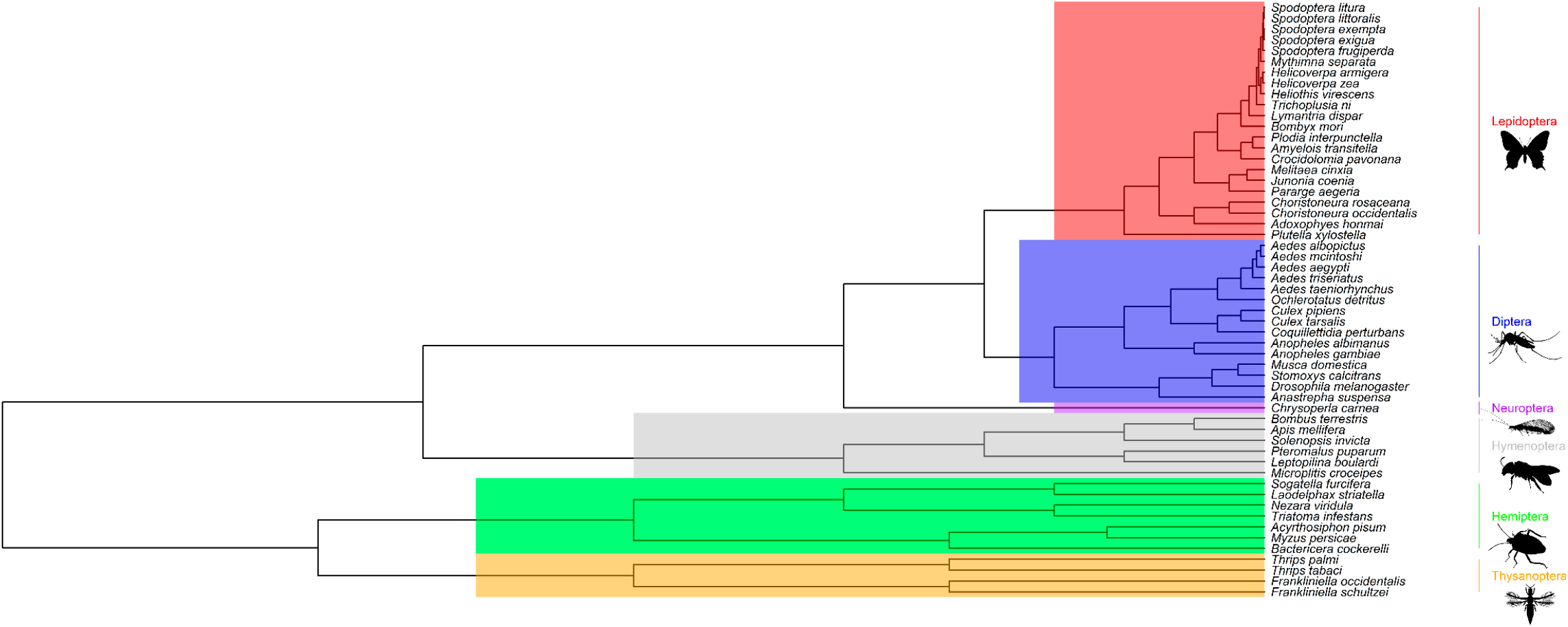
Phylogeny of insect hosts included in the meta-analysis, representing insect orders Lepidoptera, Diptera, Neuroptera, Hymenoptera, Hemiptera, and Thysanoptera.

First, we ran an overall model with random effects only, including all effect sizes (N = 1,040), then we ran separated models including the following moderators as fixed factors: host order, vector, virus type, life stage, virus inoculation, and fitness component. Because we wanted to test if the negative effect of viruses on fitness was stronger in new hosts than in natural hosts, but we did not have this information for every datapoint, we ran two separate models including only the effect sizes in which we had the information if the hosts were natural or new to the specific virus the study tested (N = 755). For this data subset, we ran an overall model including only the random effects and one model including the host type (natural or new) as moderator. We built a total of 9 models (Table S4). For each moderator we included as fixed factor, we tested if the mean effect size of each moderator level differed from zero (mean effect size), and if moderator levels differed from each other (β). All analyses were performed using the *rma*.*mv* function in the *metafor* package in R [41, 42]. We considered the effect sizes statistically significant when the 95% confidence intervals (C.I.) did not overlap zero (α = 0.05).

We used a modified version of I^2^ metric to calculate heterogeneity in the models, described in Santos and Nakagawa (2012) [44]. For each model, we quantified the total amount of heterogeneity and the percentage of contribution of each random effect for the variance in the models. We checked for potential publication bias conducting Egger’s regression [47] with the *lm* function in R [41], using the residuals from the multilevel meta-analytic models.

## Results

### Virus infection cause significant decrease on host fitness, but the decrease is higher in non-vector insects than in vectors

When analyzing all effect sizes together, mean overall effect size was negative and moderate to high, indicating that viruses cause significant reduction on host fitness (Table 1, Fig 2A).

**Table 1.**
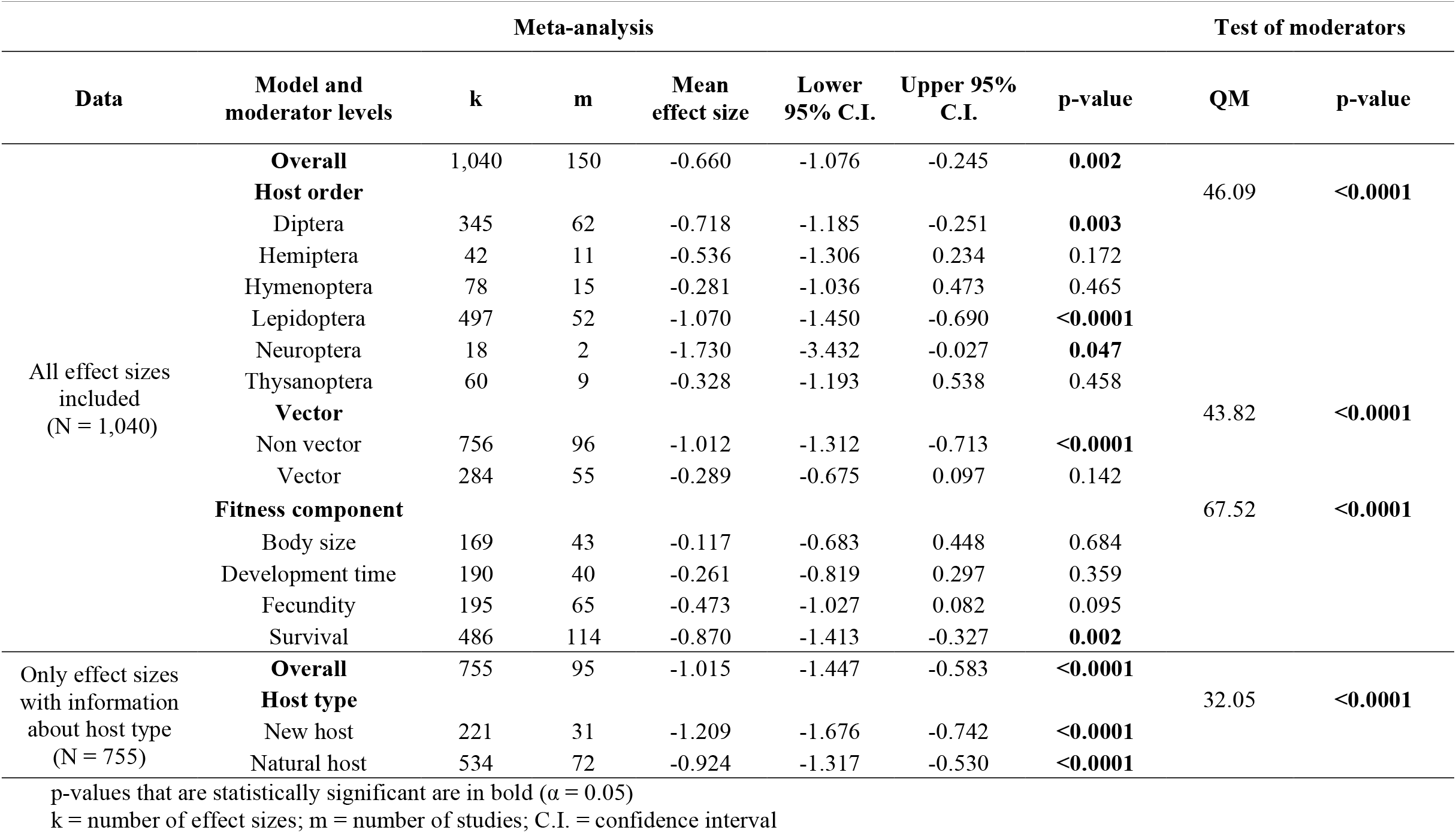
Results of the meta-analyses on the effect of virus infection on insect host and the test of moderators, including the overall models and the models including the following moderators: vector, host order, fitness component, and host type.

**Fig 2.**
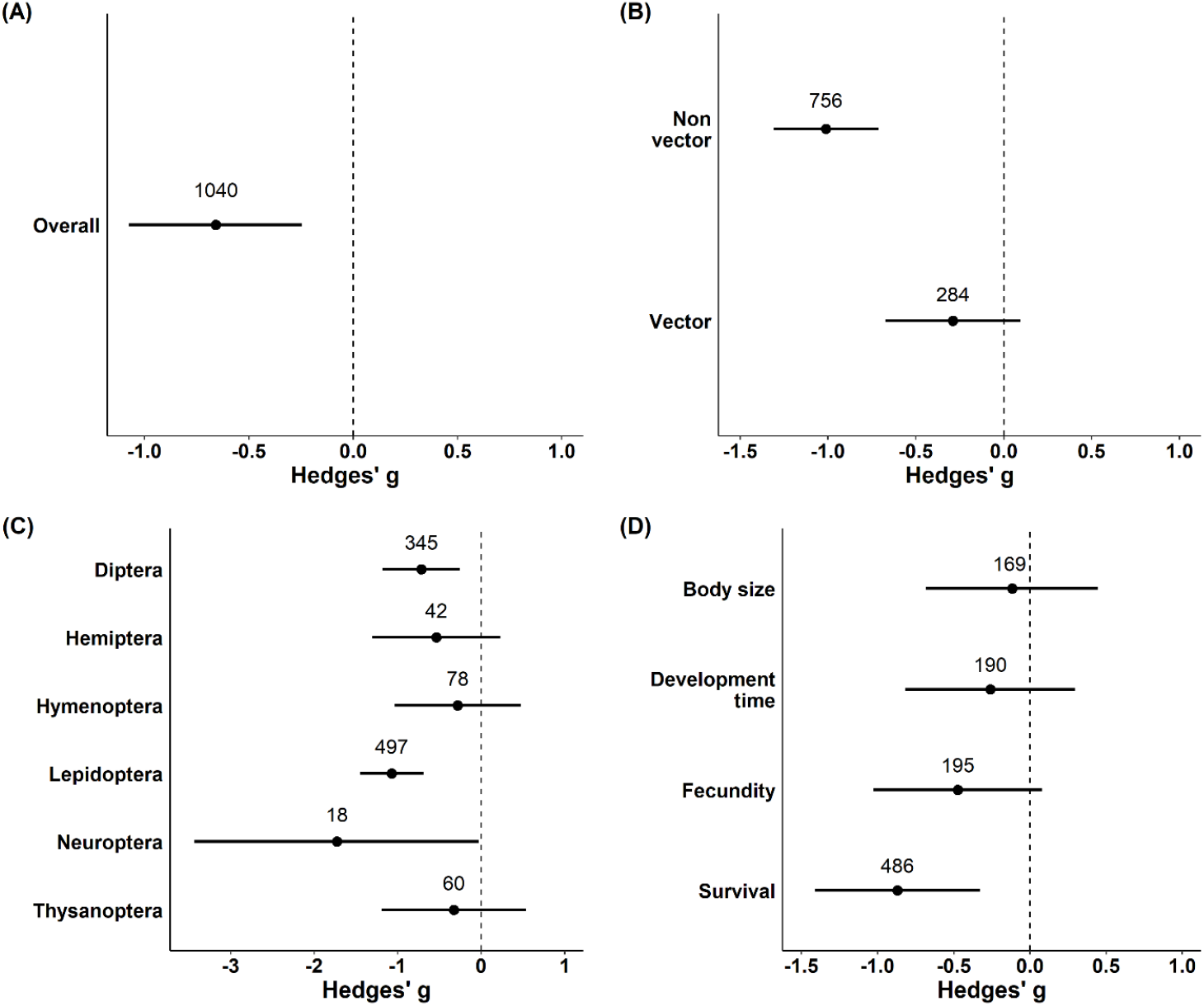
Effect of virus infection on the fitness of hosts. (A) overall model, (B) model including vector (i.e., if the insect is a vector for viruses that causes diseases in plants/animals or not) as moderator, (C) model including host order as moderator, and (D) model including fitness component as moderator. Negative values indicate a negative effect on the fitness of hosts. Therefore, negative values indicate decreased body size, fecundity, and survival, but increased development time. Points are the weighted mean effect sizes ± 95% confidence intervals. Numbers above points are the number of effect sizes.

Fitness of both vector and non-vector insects was negatively affected by viral infection, and although the upper confidence interval of vector slightly overlapped zero, we found little evidence of a positive effect due to viral infection (Table 1, Fig 2B). The magnitude of effects differed between vector and non-vector (Table S5). Non-vector mean effect size was 3.5 times larger than vector mean effect size (Table 1, Fig 2B), indicating that the reduction on fitness caused by viral infection is higher in non-vector insects than in vectors.

Fitness of hosts from the orders Diptera, Lepidoptera, and Neuroptera was negatively affected by viral infection, but not from orders Hemiptera, Hymenoptera, and Thysanoptera (Table 1, Fig 2C). However, the magnitude of effects did not differ between insect orders (Table S5).

The survival and fecundity of hosts were negatively affected by the presence of viral infection, but not body size and development time. Although the upper confidence interval of development time slightly overlapped zero, we found little evidence of a positive effect due to viral infection (Table 1, Fig 2D). The magnitude of effects differed between body size, fecundity, and survival, but not between body size and development time (Table S5).

Fitness from insects infected with DNA or RNA viruses was negatively affected (Table S6, Fig S2A). However, the magnitude of effect of DNA viruses was significantly higher than RNA viruses (Table S5). Hosts inoculated by different methods had their fitness negatively affected. Hosts naturally infected with virus had a negative mean effect size and although the upper confidence interval overlapped zero, we found little evidence of a positive effect due to viral infection (Table S6, Fig S2B). There was no difference in the magnitude of effects between the types of inoculation (Table S5). Fitness of hosts was negatively affected by viral infection at all life stages (Table S6, Fig S2C). The magnitude of effects differed only between adult and larval life stages (Table S5).

### No difference in fitness decrease caused by viral infection between new and natural hosts

When analyzing only the effect sizes from studies that had information about host type (i.e., if the insect was a new or a natural host for that specific virus), mean overall effect size was negative and high, indicating that viruses cause significant decrease on host fitness (Table 1, Fig 3A).

**Figure 3.**
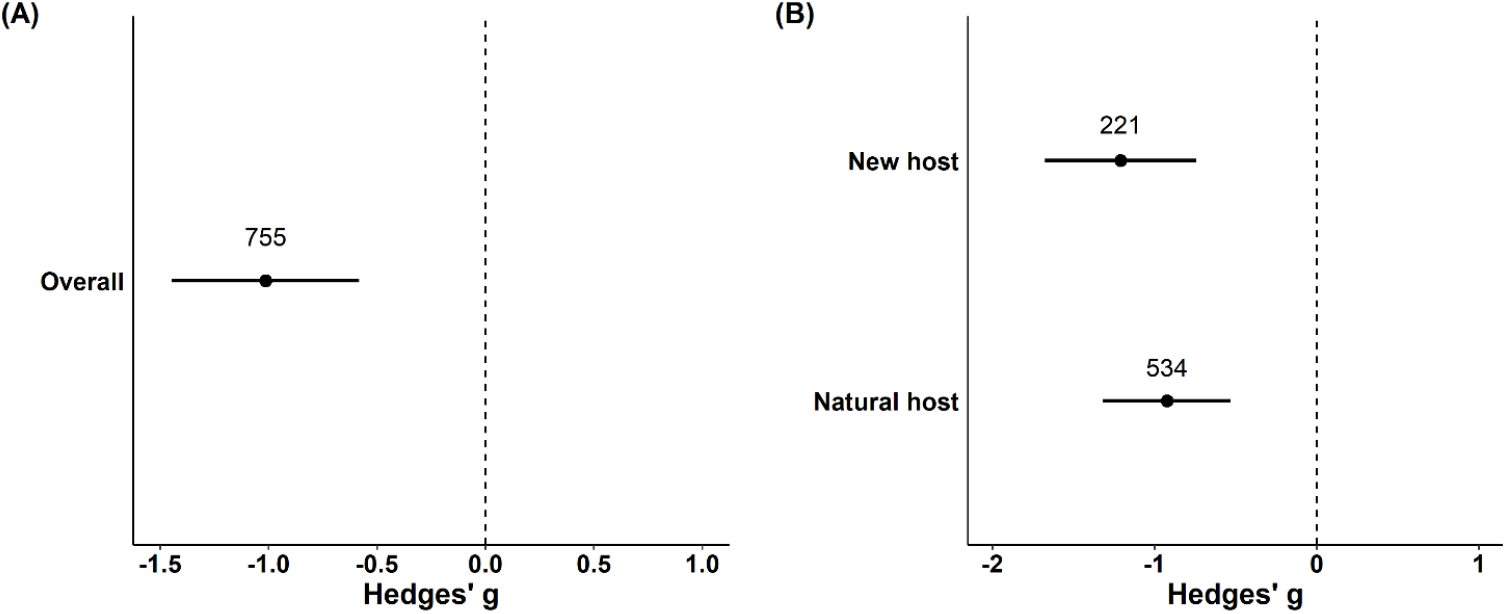
Effect of virus infection on the fitness of hosts. (A) overall model, (B) model including host type (if the insect is a natural or a new host of a specific virus) as moderator. Negative values indicate a negative effect on the fitness of hosts. Points are the weighted mean effect sizes ± 95% confidence intervals. Numbers above points are the number of effect sizes.

The fitness of both new and natural hosts was negatively affected by viral infection (Table 1, Fig 3B). Although the mean effect size of new hosts was 1.31 times larger than the mean effect size of natural hosts, the magnitude of effects did not differ between the two (Table S5). Our results suggest that the decrease in host fitness because of viral infection is only slightly higher in new hosts than in natural hosts, not being statistically significant.

### Heterogeneity and publication bias

All models presented high heterogeneity (above 95%, Table S7). In all models, the effect size ID (within study variance) explained most of the variance, followed by the study ID (between study variance), and host species. Host phylogeny explained very little of the variance in the models.

We found evidence for publication bias in four out of the nine models. The models “host order”, “vector, and “virus type” from the analysis with all effect sizes included and both models from the analysis with only effect sizes with information about host did not have evidence for publication bias. All the remaining models had evidence of publication bias (Table S8).

## Discussion

Through this meta-analysis we aimed to provide a deeper insight into the current knowledge we have on viral infection in insects, quantifying the extent of the harm that viruses can cause on the fitness of hosts, and the possible factors influencing the dynamic of insect-virus interaction. Here, we collected data from 150 studies, totalizing 1,040 effect sizes from 90 different study systems. Our results revealed that overall, viruses cause strong harmful effects on hosts by decreasing their fitness, especially their survival.

Our results show that, in general, viruses cause significant decrease on host fitness. Many studies show that most known viruses cause a great reduction in host fitness, and a meta-analysis on effects of pathogens on insect pollinators and herbivores found that viruses cause great reduction on host fitness [48], which is in accordance with our results. However, although viruses are the most abundant microorganisms in nature [49], we still know little about their diversity and biology, and most well studied viruses are pathogenic viruses that have been successfully isolated from hosts that presented changes in their phenotype because of viral infection [18, 19]. With the advances in molecular biology, recent metagenomic studies showed that wild insect populations such as flies and honeybees can be infected by up to 30 different viruses [10, 21], but only a small portion of these viruses have been isolated and studied. Hence, the effects of most recent-discovered viruses on host fitness remain largely unknown as well as the nature of these interactions, which can range from parasitic to mutualistic relationships. However, these same metagenomic studies showed that natural insect populations are also infected by pathogenic viruses that have been previously isolated and can cause high mortality rates and other adverse effects on host fitness, such as reduced fecundity. For example, Gebremedhn et al. (2020) [21] found 30 different viruses infecting honeybee colonies in apiaries, including deformed wing virus, that reduces the lifespan of bees and causes colonies to collapse [50]. Cogni et al. (2021) [10] found 30 different viruses infecting a single *D. melanogaster* population, including Kallithea virus that causes reduction in fecundity and high mortality rates in males [51], and La Jolla virus (LJV). LJV causes high mortality in different *Drosophila* species, including *D. melanogaster* and *D. suzukii* [52, 53]. Moreover, a recent study showed that five naturally occurring viruses of *D. melanogaster* can reduce fly longevity and a few of these viruses can also reduce fly fecundity [54]. Therefore, many naturally occurring viruses can act as strong selective agents for host adaptation against infection, such as the evolution of inherent immune defense traits or the association with defensive symbionts such as *Wolbachia*.

We found that the level of negative impact that viruses cause on the fitness of their hosts varied between insect orders. Hosts from orders Diptera, Lepidoptera and Neuroptera suffered a significant reduction in their fitness when infected with virus. Lepidoptera is a group that has been largely studied given its importance to the agriculture field, as many species are crop pests [17]. Most studies on virus infecting Lepidoptera are strict to the family Baculoviridae since these DNA viruses have been successfully used as biopesticide [17, 55]. This can explain the strong negative effect that we found on the fitness of lepidopterans, since Baculoviruses can cause high mortality rates in caterpillars [17, 56]. Diptera is a group of great importance to the field of host-virus interaction as it includes two major groups of hosts that are studied: *Drosophila* and mosquitoes. *Drosophila* species have been used as models to study adaptations to resist pathogens and the evolution of host-virus interaction in general, including studies that show that bacterial symbiont *Wolbachia* can protect flies against viral infection [3, 5, 8, 9, 57]. Most of these studies are on RNA viruses from families Picornaviridae and Dicistroviridae, such as Nora and DCV, respectively [9, 58, 59]. Mosquitoes have been largely studied since they are vectors of insect-borne diseases such as Dengue and Chikungunya, and most studies on the interaction between mosquitoes-viruses are strict to Arboviruses that cause these diseases [24, 25, 29]. Most studies on *Drosophila* were done with RNA viruses that cause high mortality rates [60], and studies on mosquitoes shows that Arboviruses can cause reduction in their fitness despite them being vectors [61, 62]. Insects from orders Hymenoptera, Hemiptera and Thysanoptera also had a negative effect on their fitness because of viral infection. However, this effect is lower and not as clear as seen in the other host orders. Most of our data on Hemiptera and Thysanoptera come from hosts that are vectors of plant viruses, belonging to different virus families. Although viruses can cause reduction in the fitness of their insect vectors, there are cases in which viruses facilitate their spread by increasing fecundity of their hosts. For example, a study with *Frankliniella fusca* (Thysanoptera) showed that insects exposed to tomato spotted wilt virus infected plants had higher fecundity than insects exposed to uninfected plants [63]. Hymenoptera hosts have some interesting cases in which viruses become mutualists of parasitoid wasps. For example, some wasps deposit their eggs inside caterpillars together with polydnavirus virions, preventing egg encapsulation and increasing the survival rate of parasitoids [22, 64]. Cases of viruses that increase the fitness of their vectors or are mutualists can help explain the results we found for orders Hymenoptera, Hemiptera and Thysanoptera. Therefore, the extent to which viruses negatively impact host fitness varies across different insect orders and is influenced by the specific biological traits of the viruses infecting these groups, which includes whether the viruses have coevolved with their hosts to establish parasitic, commensal, or mutualistic relationships.

However, most viruses discovered and studied so far established a parasitic relationship with their hosts and only a few mutualistic relationships between viruses and insects have been described.

Our results show that non-vector insects had their fitness more significantly reduced than vectors. The vector-pathogen dynamic is a particular case as viruses can have two distinct hosts. Taking the case of pathogens and mosquitos for example, usually, mosquito-borne pathogens are generalists that infect multiple insect vectors and vertebrate species before evolving to specialize in infecting humans together with their vectors [65]. Hence, pathogens such as Arboviruses for example, have been coevolving with their vectors for a long time before becoming pathogens of humans. Mosquitoes have complex antiviral defenses that limit viral replication, but their immune system often may not be able to eliminate the virus completely [66], most likely because Arboviruses have evolved adaptations to fight the resistance mechanism of hosts [67], allowing viral infection to spread in the mosquito system and favoring vector competence (i.e., the ability of transmitting infectious agents) [68, 69]. Moreover, because infections with Arboviruses are usually not lethal to mosquitoes, they can carry and transmit viruses for extended periods of time, making them efficient vectors [66, 69]. Therefore, because viruses depend on the success of their vector to complete their cycle and be transmitted to their next host, these viruses might have evolved to be less virulent and cause less damage on the fitness of vectors in comparison to non-vector insects, corroborating the results we found.

Regarding the origin of the virus, our results show that new hosts have slightly higher, but non-significant, fitness decrease due to viral infection in comparison to natural hosts, suggesting a certain degree of similarity in host susceptibility and the general impact of the virus across host species. The susceptibility of hosts and virulence can vary at an evolutionary scale, usually with recent host-virus associations having higher susceptibility and virulence [31, 32], therefore causing greater negative impact on the fitness of hosts. However, our results suggest that new hosts may not always experience disproportionately severe fitness costs, as commonly assumed. In general, viruses must overcome the immune defense of hosts to successfully establish itself into a new host [70]. Because viruses – especially RNA viruses – have high mutation rates, they have higher capacity to accumulate the necessary mutations to adapt to new hosts in comparison to other pathogens [32, 71]. The phylogenetic relationship between hosts is also important when it comes to new host-virus associations, since closely related species may have similar levels of susceptibility and have similar environments for the virus to adapt [31, 32, 72]. For example, Longdon et al. (2011) [72] infected 51 different species of *Drosophila* with three strains of Sigma virus and found that new host species that were more closely related to the original host had higher viral replication. Therefore, while it is expected that new hosts have greater loss in their fitness than natural hosts that are already well adapted to the virus, closely related host species might suffer similar levels of fitness decrease, which may explain our results.

Besides traits of insect hosts, such as if they are vectors or non-vectors of viruses that causes diseases or if they are new or natural hosts of the virus, other biotic and abiotic factors can influence the outcome of the host-virus association [73]. The presence of bacterial symbionts is an important biotic factor that directly affects the interaction between hosts and viruses. Many bacterial symbionts can protect their hosts against natural enemies such as fungi, parasitoid wasps, nematodes, and viruses [74, 75]. *Wolbachia* is the most common bacterial symbiont, found infecting around 50% of all sampled terrestrial arthropod species, including many insect species from different orders [76]. *Wolbachia* is known for protecting flies and mosquitoes hosts from viral infection and many studies have shown that *Wolbachia* can increase the fitness of their hosts by increasing host survival when they are infected with virus [8, 9, 14, 77]. Here, we found that viruses cause a significant decrease in host fitness, especially host survival. Therefore, although interactions with symbionts have a cost associated, this cost can be overcome and compensated by *Wolbachia*’s mutualistic effect of protecting their hosts by increasing their fitness when hosts are infected with viruses. Moreover, the presence of virus infection in hosts may act as selective pressure for *Wolbachia* to spread in wild insect populations.

In conclusion, the current knowledge we have on viral infection in insects shows that most host-virus interactions evolved to become parasitic, with viruses exerting significant decrease on the fitness of hosts. Moreover, host traits, such as whether the insect is a vector of viruses that cause diseases in humans or plants, can directly affect the level of negative impact that viruses have on host fitness. However, as recent metagenomic studies have shown, we still lack knowledge about viral diversity in wild insect populations, and only a small fraction of this diversity has been described. New viruses are continually being discovered, and their effects on host fitness must be investigated to improve our current knowledge. Addressing these questions is crucial and future studies should focus on exploring virus diversity and whether most of these viruses have coevolved with their hosts to establish parasitic, commensal, or mutualistic relationships. However, although much remains to be investigated, recent studies have shown that recent discovered viruses found infecting wild insect populations can cause substantial reduction on host fitness, which is in accordance with our results – most host-virus associations evolved to become parasitic. In this context, harboring symbionts that confer antiviral protection by enhancing their fitness, such as *Wolbachia*, can be highly advantageous for hosts. Conversely, *Wolbachia* can also benefit from the presence of viruses, facilitating their spread in host populations by offering antiviral protection for hosts.

## Acknowledgments

We would like to thank Leonardo M. Servino for the help with insect phylogeny and Figure 1. This study was financed by Conselho Nacional de Desenvolvimento Científico e Tecnológico (CNPq; 307015/2015-7 and 307447/2018-9) and the São Paulo Research Foundation (FAPESP), Brasil. FAPESP processes numbers: 2019/03997-2 and 2021/13166-0 (Cássia S. Cesar); 2013/25991-0 and 2021/06874-9 (Rodrigo Cogni).

## Author contributions

Conceptualization: Cássia S. Cesar, Rodrigo Cogni

Data collection: Cássia S. Cesar, Vitória H. Miranda, Elielson R. Silveira, Thiago A. Oliveira

Analysis: Cássia S. Cesar

Writing: Cássia S. Cesar, Rodrigo Cogni

## Data availability

All data and R script will be available at GitHub.

