## Supplementary information for "Virus infection significantly decreases insect fitness: a meta-analysis"

**List of papers included in the meta-analysis with their respective ID in the raw data tables.** Papers with * were excluded during the analysis as described at the Methods section of the manuscript.

10. Martin E, Moutailler S, Madec Y, Failloux AB. 2010 Differential responses of the mosquito *Aedes albopictus* from the Indian Ocean region to two chikungunya isolates. *BMC Ecol* **10**, 8. (doi:10.1186/1472-6785-10-8) (P153)
11. Ciota AT, Styer LM, Meola MA, Kramer LD. 2011 The costs of infection and resistance as determinants of West Nile virus susceptibility in *Culex* mosquitoes. *BMC Ecol* **11**, 23. (doi:10.1186/1472-6785-11-23) (P154)
12. Mahiba Helen S, Sudheendrakumar V V., Sajeev T V., Deshmuk RA. 2012 Biosafety of crude and formulated *Hyblaea puera* (Cramer) (Lepidoptera: Hyblaeidae), Nucleopolyherovirus (HpNPV) against silkworm *Bombyx mori* (L.), Indian mynah, *Acridotherus tristis* (Linn.) and cell lines. *J Biopesticides* **5**, 201-207. (P146)
13. Rodriguez-Andres J *et al.* 2012 Phenoloxidase activity acts as a mosquito innate immune response against infection with Semliki Forest Virus. *PLoS Pathog* **8**, e1002977. (doi:10.1371/journal.ppat.1002977) (P147)
14. Desai SD, Eu YJ, Whyard S, Currie RW. 2012 Reduction in deformed wing virus infection in larval and adult honey bees (*Apis mellifera* L.) by double-stranded RNA ingestion. *Insect Mol Biol* **21**, 446-455. (doi:10.1111/j.1365-2583.2012.01150.x) (P148)
15. Azzami K, Ritter W, Tautz J, Beier H. 2012 Infection of honeybees with acute bee paralysis virus does not trigger humoral or cellular immune responses. *Arch Virol* **157**, 689-702. (doi:10.1007/s00705-012-1223-0) (P149)
16. Arnold PA, Johnson KN, White CR. 2013 Physiological and metabolic consequences of viral infection in *Drosophila melanogaster*. *J Exp Biol* **216**, 3350-3357. (doi:10.1242/jeb.088138) (P144)
17. Subbaiah E V. *et al.* 2013 Engineering silkworms for resistance to baculovirus through multigene RNA interference. *Genetics* **193**, 63-75. (doi:10.1534/genetics.112.144402) (P145)
18. Retschnig G, Williams GR, Mehmann MM, Yañez O, De Miranda JR, Neumann P. 2014 Sex-specific differences in pathogen susceptibility in honey bees (*Apis mellifera*). *PLoS One* **9**, e0085261. (doi:10.1371/journal.pone.0085261) (P162)
19. Xu K, Li F, Ma L, Wang B, Zhang H, Ni M, Hong F, Shen W, Li B. 2015 Mechanism of enhanced *Bombyx mori* nucleopolyhedrovirus-resistance by titanium dioxide nanoparticles in silkworm. *PLoS One* **10**, e0118222. (doi:10.1371/journal.pone.0118222) (P163)
20. Stevanovic AL, Johnson KN. 2015 Infectivity of *Drosophila* C virus following oral delivery in drosophila larvae. *J Gen Virol* **96**, 1490-1496. (doi:10.1099/vir.0.000068) (P164)
21. Zhang S, Wu F, Li Z, Lu Z, Zhang X, Zhang Q, Liu X. 2015 Effects of nucleopolyhedrovirus infection on the development of *Helicoverpa armigera* (Lepidoptera: Noctuidae) and

expression of its 20-hydroxyecdysone-and juvenile hormone-related genes. *Fla Entomol* **98**, 682-689. (doi:10.1653/024.098.0243) (P165)

evaluation of vector status of *Frankliniella tritici* and *Frankliniella fusca*. *J Econ Entomol* **109**, 1979-1987. (doi:10.1093/jee/tow145) (P008)

### Supplementary figures

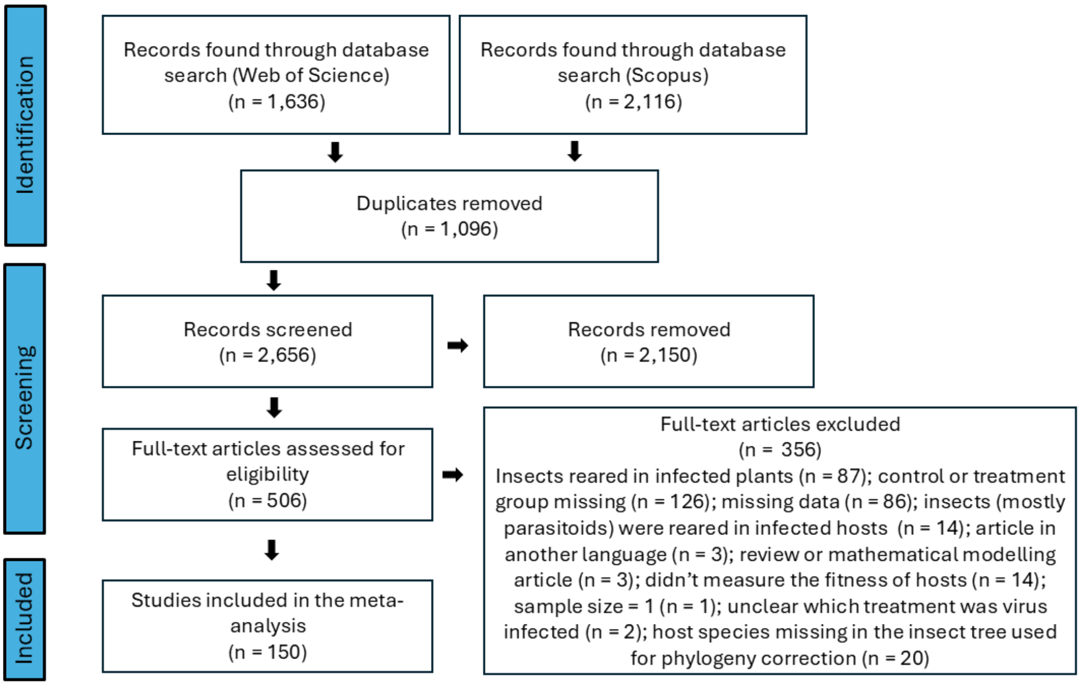

**Figure S1**. Full screening process of inclusion and exclusion of studies following the PRISMA protocol.

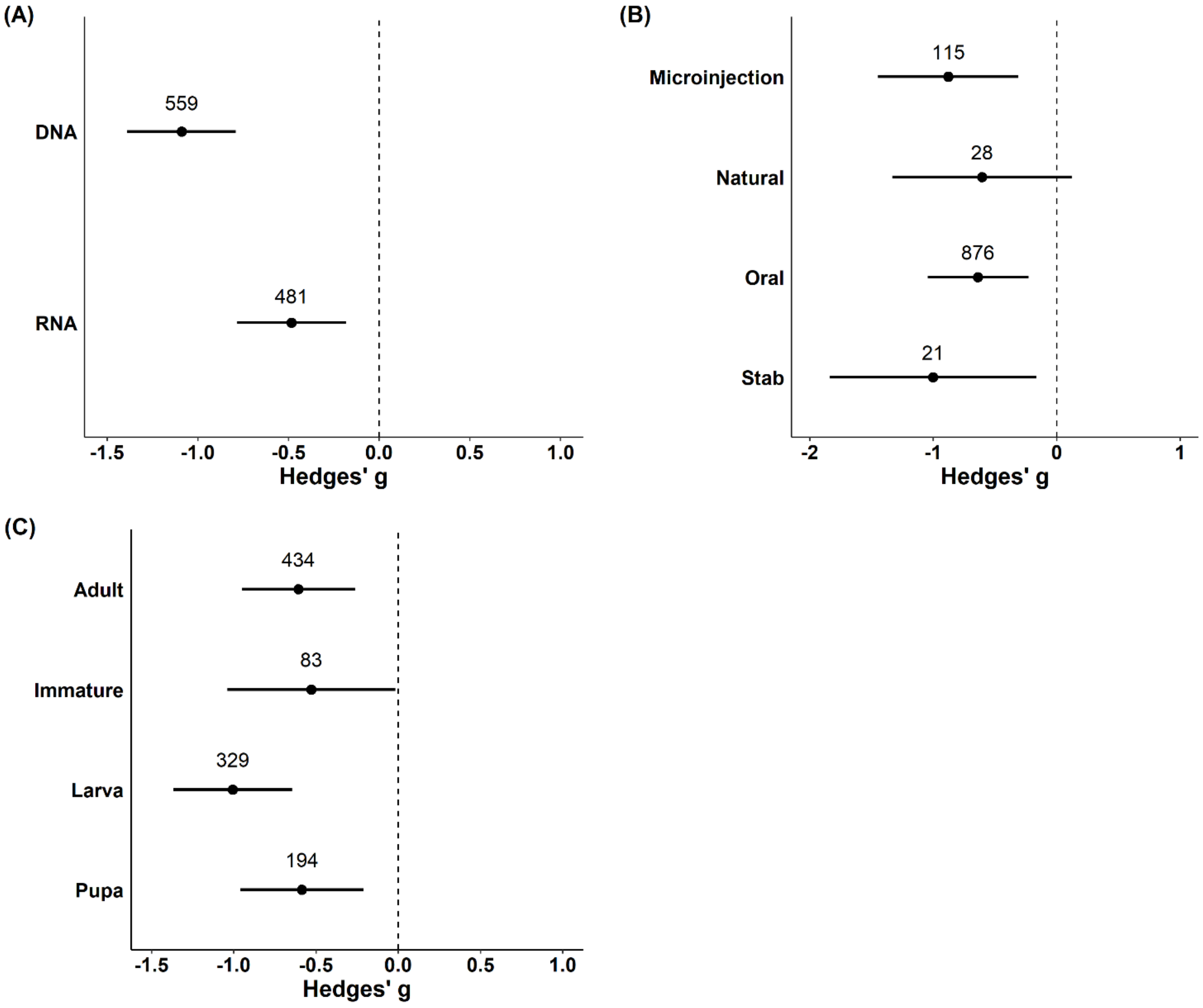

**Figure S2.** Effect of viruses on the fitness of hosts. (a) model including virus type (if it is a DNA or RNA virus) as moderator, (b) model including virus inoculation (method used to inoculate the virus in hosts) as moderator, and (c) model including life stage (life stage of the host in which the viral effect was measured) as moderator. Negative values indicate a negative effect on the fitness of hosts. Points are the weighted mean effect sizes ± 95% confidence intervals. Numbers above points are the number of effect sizes.

### Supplementary tables

**Table S1.** Study systems included in the meta-analysis.

| Host species | Host order | Virus family |
|---|---|---|
| *Aedes aegypti* | Diptera | Bunyaviridae |
| *Aedes aegypti* | Diptera | Flaviviridae |
| *Aedes aegypti* | Diptera | Iridoviridae |
| *Aedes aegypti* | Diptera | Parvoviridae |
| *Aedes aegypti* | Diptera | Togaviridae |
| *Aedes albopictus* | Diptera | Flaviviridae |
| *Aedes albopictus* | Diptera | Togaviridae |
| *Aedes mcintoshi* | Diptera | Bunyaviridae |
| *Aedes taeniorhynchus* | Diptera | Bunyaviridae |
| *Aedes taeniorhynchus* | Diptera | Flaviviridae |
| *Aedes taeniorhynchus* | Diptera | Togaviridae |
| *Aedes triseriatus* | Diptera | Bunyaviridae |
| *Anastrepha suspensa* | Diptera | Poxviridae |
| *Anopheles albimanus* | Diptera | Bunyaviridae |
| *Anopheles gambiae* | Diptera | Parvoviridae |
| *Anopheles gambiae* | Diptera | Togaviridae |
| *Coquillettidia perturbans* | Diptera | Togaviridae |
| *Culex pipiens* | Diptera | Bunyaviridae |
| *Culex pipiens* | Diptera | Flaviviridae |
| *Culex tarsalis* | Diptera | Flaviviridae |
| *Culex tarsalis* | Diptera | Togaviridae |
| *Drosophila melanogaster* | Diptera | Artoviridae |
| *Drosophila melanogaster* | Diptera | Dicistroviridae |
| *Drosophila melanogaster* | Diptera | Iridoviridae |
| *Drosophila melanogaster* | Diptera | Nodaviridae |
| *Drosophila melanogaster* | Diptera | Nudiviridae |
| *Drosophila melanogaster* | Diptera | Picornaviridae |
| *Drosophila melanogaster* | Diptera | Rhabdoviridae |
| *Drosophila melanogaster* | Diptera | Togaviridae |
| *Musca domestica* | Diptera | Hytrosaviridae |
| *Ochlerotatus detritus* | Diptera | Bunyaviridae |
| *Stomoxys calcitrans* | Diptera | Hytrosaviridae |
| *Acyrthosiphon pisum* | Hemiptera | Picornaviridae |
| *Bactericera cockerelli* | Hemiptera | Potyviridae |
| *Laodelphax striatella* | Hemiptera | Phenuiviridae |
| *Laodelphax striatella* | Hemiptera | Rhabdoviridae |
| *Myzus persicae* | Hemiptera | Luteoviridae |
| *Nezara viridula* | Hemiptera | Picornaviridae + Totiviridae |
| *Sogatella furcifera* | Hemiptera | Reoviridae |
| *Triatoma infestans* | Hemiptera | Picornaviridae |

| | | |
|---|---|---|
| *Apis mellifera* | Hymenoptera | Dicistroviridae |
| *Apis mellifera* | Hymenoptera | Iflaviridae |
| *Apis mellifera* | Hymenoptera | Nodaviridae |
| *Apis mellifera* | Hymenoptera | Parvoviridae |
| *Bombus terrestris* | Hymenoptera | Iflaviridae |
| *Leptopilina boulardi* | Hymenoptera | Filamentoviridae |
| *Leptopilina boulardi* | Hymenoptera | Totiviridae |
| *Microplitis croceipes* | Hymenoptera | Baculoviridae |
| *Pteromalus puparum* | Hymenoptera | Nyamiviridae |
| *Solenopsis invicta* | Hymenoptera | Togaviridae |
| *Adoxophyes honmai* | Lepidoptera | Baculoviridae |
| *Adoxophyes honmai* | Lepidoptera | Poxviridae |
| *Adoxophyes honmai* | Lepidoptera | Poxviridae + Baculoviridae |
| *Amyelois transitella* | Lepidoptera | Closteroviridae + Calciviridae |
| *Bombyx mori* | Lepidoptera | Baculoviridae |
| *Choristoneura occidentalis* | Lepidoptera | Baculoviridae |
| *Choristoneura rosaceana* | Lepidoptera | Baculoviridae |
| *Crocidolomia pavonana* | Lepidoptera | Ascoviridae |
| *Helicoverpa armigera* | Lepidoptera | Ascoviridae |
| *Helicoverpa armigera* | Lepidoptera | Baculoviridae |
| *Helicoverpa armigera* | Lepidoptera | Iflaviridae |
| *Helicoverpa armigera* | Lepidoptera | Parvoviridae |
| *Helicoverpa zea* | Lepidoptera | Baculoviridae |
| *Heliothis virescens* | Lepidoptera | Baculoviridae |
| *Junonia coenia* | Lepidoptera | Dicistroviridae |
| *Junonia coenia* | Lepidoptera | Parvoviridae |
| *Lymantria dispar* | Lepidoptera | Baculoviridae |
| *Melitaea cinxia* | Lepidoptera | Baculoviridae |
| *Mythimna separata* | Lepidoptera | Baculoviridae |
| *Pararge aegeria* | Lepidoptera | Baculoviridae |
| *Plodia interpunctella* | Lepidoptera | Baculoviridae |
| *Plutella xylostella* | Lepidoptera | Ascoviridae |
| *Plutella xylostella* | Lepidoptera | Baculoviridae |
| *Spodoptera exempta* | Lepidoptera | Baculoviridae |
| *Spodoptera exempta* | Lepidoptera | Partitiviridae |
| *Spodoptera exigua* | Lepidoptera | Ascoviridae |
| *Spodoptera exigua* | Lepidoptera | Baculoviridae |
| *Spodoptera exigua* | Lepidoptera | Iflaviridae |
| *Spodoptera frugiperda* | Lepidoptera | Baculoviridae |
| *Spodoptera frugiperda* | Lepidoptera | Iridoviridae |
| *Spodoptera frugiperda* | Lepidoptera | Partitiviridae |
| *Spodoptera littoralis* | Lepidoptera | Baculoviridae |
| *Spodoptera litura* | Lepidoptera | Ascoviridae |
| *Spodoptera litura* | Lepidoptera | Baculoviridae |

| | | |
|---|---|---|
| *Trichoplusia ni* | Lepidoptera | Baculoviridae |
| *Chrysoperla carnea* | Neuroptera | Baculoviridae |
| *Frankliniella occidentalis* | Thysanoptera | Tospoviridae |
| *Frankliniella schultzei* | Thysanoptera | Tospoviridae |
| *Thrips palmi* | Thysanoptera | Tospoviridae |
| *Thrips tabaci* | Thysanoptera | Tospoviridae |

**Table S2.** List of all fitness measures extracted from the studies to calculate the mean effect sizes. Measures are grouped in four broad categories of fitness component: body size, development time, fecundity, and survival.

| **Category** | **Fitness measure** |
|---|---|
| Body size | Wing length, head width, cocoon length, weight |
| Development time | Development time (Time taken to develop from one life stage into another. Examples: larva to pupa, pupa to adult, larva to adult, nymph to adult) |
| Fecundity | Fecundity (number of eggs), fertility (number of ovarioles or oocytes), egg hatch (% or number of eggs that hatched), number of adult offspring |
| Survival | Longevity (mean number of days alive during a life stage), lifespan (mean number of days alive from hatch until death), survival (% or number of individuals alive), emergence rate (% or number of pupae that hatched) |

**Table S3.** Variables collected from the original studies to be used as moderators in the models.

| Moderator | Moderator level |
|---|---|
| Host order | Diptera |
| | Hemiptera |
| | Hymenoptera |
| | Lepidoptera |
| | Neuroptera |
| | Thysanoptera |
| Fitness component | Body size |
| | Development time |
| | Fecundity |
| | Survival |
| Vector | Non vector |
| | Vector |
| Host type | New host |
| | Natural host |
| Virus type | DNA |
| | RNA |
| Life stage | Adult |
| | Embryo |
| | Immature (insect nymphs or when authors did not separate larval and pupal stages) |
| | Larva |
| | Pupa |
| Virus inoculation | Microinjection |
| | Natural (when hosts were infected with viruses that are vertically transmitted or were collected from wild populations and already infected with virus) |
| | Oral (hosts fed with viral solution or infected food) |
| | Stab (hosts stabbed with a pin dipped in viral solution) |

**Table S4.** Multilevel meta-analytical models performed in this study, specifying their random and fixed factors (moderators) and moderator levels.

| Data | Model | Random factors | Fixed factors (moderators) | Moderator levels |
|---|---|---|---|---|
| All effect sizes included (N = 1,040) | Overall | Study ID + effect size ID + Host phylogeny + Host species | - | - |
| | Host order | Study ID + effect size ID + Host phylogeny + Host species | Host order | Diptera<br>Hemiptera<br>Hymenoptera<br>Lepidoptera<br>Neuroptera<br>Thysanoptera |
| | Vector | Study ID + effect size ID + Host phylogeny + Host species | Vector | Non vector<br>Vector |
| | Fitness component | Study ID + effect size ID + Host phylogeny + Host species | Fitness component | Body size<br>Development time<br>Fecundity<br>Survival |
| | Virus type | Study ID + effect size ID + Host phylogeny + Host species | Virus type | DNA<br>RNA |
| | Life stage | Study ID + effect size ID + Host phylogeny + Host species | Life stage | Adult<br>Immature<br>Larva<br>Pupa |
| | Virus inoculation | Study ID + effect size ID + Host phylogeny + Host species | Virus inoculation | Microinjection<br>Natural<br>Oral<br>Stab |
| Only effect sizes with information about host type (N = 755) | Overall | Study ID + effect size ID + Host phylogeny + Host species | - | - |
| | Host type | Study ID + effect size ID + Host phylogeny + Host species | Host type | New host<br>Natural host |

**Table S5.** Results of the meta-analyses on the effect of virus infection on insect hosts. Models including the first moderator level as intercept to test if moderator levels differed from one another.

| | Meta-analysis | | | | | Test of moderators | |
|---|---|---|---|---|---|---|---|
| **Data** | **Model and moderator levels** | **Slope (β)** | **Lower 95% C.I.** | **Upper 95% C.I.** | **p-value** | **QM** | **p-value** |
| All effect sizes included (N = 1,040) | **Host order** | | | | | 6.55 | 0.255 |
| | Intercept (Diptera) | -0.718 | -1.185 | -0.251 | **0.003** | | |
| | Hemiptera | 0.181 | -0.719 | 1.082 | 0.693 | | |
| | Hymenoptera | 0.436 | -0.451 | 1.324 | 0.335 | | |
| | Lepidoptera | -0.352 | -0.954 | 0.250 | 0.252 | | |
| | Neuroptera | -1.012 | -2.777 | 0.754 | 0.261 | | |
| | Thysanoptera | 0.390 | -0.594 | 1.374 | 0.437 | | |
| | **Vector** | | | | | 10.81 | **0.001** |
| | Intercept (Non vector) | -1.012 | -1.312 | -0.713 | **<0.0001** | | |
| | Vector | 0.723 | 0.292 | 1.155 | **0.001** | | |
| | **Fitness component** | | | | | 62.75 | **<0.0001** |
| | Intercept (Body size) | -0.117 | -0.683 | 0.448 | 0.684 | | |
| | Development time | -0.144 | -0.363 | 0.076 | 0.199 | | |
| | Fecundity | -0.355 | -0.597 | -0.114 | **0.004** | | |
| | Survival | -0.752 | -0.965 | -0.540 | **<0.0001** | | |
| | **Virus type** | | | | | 10.53 | **0.001** |
| | Intercept (DNA) | -1.090 | -1.389 | -0.791 | **<0.0001** | | |
| | RNA | 0.607 | 0.241 | 0.974 | **0.001** | | |
| | **Virus inoculation** | | | | | 1.81 | 0.613 |
| | Intercept (Microinjection) | -0.879 | -1.448 | -0.310 | **0.002** | | |
| | Natural | 0.273 | -0.435 | 0.982 | 0.449 | | |
| | Oral | 0.240 | -0.225 | 0.706 | 0.311 | | |
| | Stab | -0.124 | -0.946 | 0.699 | 0.768 | | |
| | **Life stage** | | | | | 20.23 | **0.0002** |
| | Intercept (Adult) | -0.607 | -0.950 | -0.263 | **0.0005** | | |
| | Immature | 0.078 | -0.332 | 0.488 | 0.709 | | |
| | Larva | -0.400 | -0.624 | -0.176 | **0.0004** | | |

| | | | | | | | |
|---|---|---|---|---|---|---|---|
| | Pupa | 0.020 | -0.213 | 0.253 | 0.865 | | |
| Only effect sizes with information about host type (N = 755) | **Host type** | | | | | 1.56 | **0.211** |
| | Intercept (Natural host) | -0.924 | -1.317 | -0.530 | **<0.0001** | | |
| | New host | -0.285 | -0.733 | 0.162 | 0.211 | | |

p-values that are statistically significant are in bold (α = 0.05)

C.I. = confidence interval

**Table S6.** Results of the meta-analyses on the effect of virus infection on insect hosts and the test of moderators for the following moderators: virus type, virus inoculation and life stage.

| | Meta-analysis | | | | | | | Test of moderators | |
|---|---|---|---|---|---|---|---|---|---|
| Data | Model and moderator levels | k | m | Mean effect size | Lower 95% C.I. | Upper 95%C.I. | p-value | QM | p-value |
| All effect sizes included (N = 1,040) | **Virus type** | | | | | | | 53.06 | **<0.0001** |
| | DNA | 559 | 63 | -1.090 | -1.389 | -0.791 | **<0.0001** | | |
| | RNA | 481 | 88 | -0.483 | -0.783 | -0.183 | **0.001** | | |
| | **Virus inoculation** | | | | | | | 12.49 | **0.014** |
| | Microinjection | 115 | 20 | -0.879 | -1.448 | -0.310 | **0.002** | | |
| | Natural | 28 | 8 | -0.606 | -1.334 | 0.122 | 0.103 | | |
| | Oral | 876 | 121 | -0.639 | -1.047 | -0.230 | **0.002** | | |
| | Stab | 21 | 6 | -1.003 | -1.841 | -0.164 | **0.019** | | |
| | **Life stage** | | | | | | | 37.97 | **<0.0001** |
| | Adult | 434 | 104 | -0.607 | -0.950 | -0.263 | **0.001** | | |
| | Immature | 83 | 16 | -0.529 | -1.039 | -0.018 | **0.042** | | |
| | Larva | 329 | 64 | -1.007 | -1.369 | -0.645 | **<0.0001** | | |
| | Pupa | 194 | 34 | -0.586 | -0.961 | -0.212 | **0.002** | | |

p-values that are statistically significant are in bold (α =0.05)
k = number of effect sizes; m = number of studies; C.I. = confidence interval

**Table S7.** Results of the heterogeneity test ($I^2$) for each of the nine meta-analytical models. For each model, we quantified the total amount of heterogeneity ($I^2$) and the percentage of contribution of each random effect for the variance in the models: study, effect size, host phylogeny and host species.

| Data | Model | $I^2$ total (%) | $I^2$ Study (%) | $I^2$ Effect size (%) | $I^2$ Host phylogeny (%) | $I^2$ Host species (%) |
|---|---|---|---|---|---|---|
| All effect sizes included (N = 1,040) | Overall | 98.59 | 26.25 | 46.00 | 4.20 | 22.14 |
| | Host order | 98.54 | 28.54 | 47.50 | 0.000076 | 22.50 |
| | Vector | 98.47 | 27.50 | 49.48 | 0.40 | 21.09 |
| | Fitness component | 98.56 | 23.74 | 43.57 | 10.46 | 20.79 |
| | Virus type | 98.47 | 26.87 | 49.76 | 0.000001 | 21.84 |
| | Virus inoculation | 98.57 | 25.89 | 46.82 | 3.76 | 22.10 |
| | Life stage | 98.51 | 25.34 | 47.97 | 1.80 | 23.40 |
| Only effect sizes with information about host type (N = 755) | Overall | 99.01 | 25.36 | 42.47 | 1.02 | 30.16 |
| | Host type | 98.96 | 26.44 | 44.46 | 0.20 | 27.86 |

**Table S8.** Summary of Egger's regression test for publication bias for each of the nine meta-analytical models. Models with p-values in bold represent the models with potential publication bias.

| Data | Model | Intercept | t | p-value |
|---|---|---|---|---|
| All effect sizes included (N = 1,040) | Overall | -0.482 | -2.685 | **0.007** |
| | Host order | -0.305 | -1.714 | 0.087 |
| | Vector | -0.332 | -1.852 | 0.064 |
| | Fitness component | -0.578 | -3.208 | **0.001** |
| | Virus type | -0.339 | -1.895 | 0.058 |
| | Virus inoculation | -0.470 | -2.616 | **0.009** |
| | Life stage | -0.415 | -2.315 | **0.020** |
| Only effect sizes with information about host type (N = 755) | Overall | -0.386 | -1.575 | 0.116 |
| | Host type | -0.396 | -1.62 | 0.106 |

p-values that are statistically significant are in bold ($\alpha$ =0.05)
